# Variations in the temperature dependence of photosynthesis among nine common tree species planted in Singapore

**DOI:** 10.1101/2024.10.19.618612

**Authors:** Li Ming Rachel Teo, Liyao Yu, Xiangzhong Luo

## Abstract

Tree planting is regarded as one of the most cost-effective ways to mitigate climate change impacts. As a part of the global tree planting movement, the tropical city-state of Singapore initiated a project to plant one million trees in 10 years. However, the environmental impacts (e.g., carbon sequestration, urban cooling) of this reforestation effort are challenging to assess as historically we lack understanding of key plant traits (i.e. photosynthetic capacity, or V_cmax_) for species in tropical Asia. Here, we conducted a comprehensive survey of key photosynthetic traits of the nine most popular tree species planted in Singapore and their short-term temperature dependence. We found a three-fold inter-species variation in their photosynthetic capacities (e.g., V_cmax_ ranges from 20 to 60 μmol m^−2^ s^−1^) as well as a large discrepancy in their temperature dependence. The photosynthesis of some species (e.g., Tembusu) is more sensitive to rising temperature than others (e.g., Mangroves) while a commonly-used photosynthesis model consistently overestimates photosynthesis but underestimates many species’ temperature sensitivities. The species with smaller temperature sensitivity of photosynthesis is likely related to their stronger ability to adjust leaf temperature via transpiration. Our study provides critical information on the temperature-dependence of photosynthesis of tree species in tropical Asia, which can be used to guide reforestation in Singapore and broadly in Southeast Asia.

## Introduction

Tree planting is considered the most cost-effective way to mitigate climate change impacts (Griscom et al. 2017, Lewis et al. 2019), as trees remove CO_2_ from the atmosphere through photosynthesis and store carbon in biomass. Many initiatives, such as the global One Billion Tree Project (Goymer 2018) and the One Million Trees movement in Singapore (NParks, 2024), are utilizing the carbon uptake of trees to offset part of the anthropogenic CO_2_ emissions. However, the photosynthetic capacities of tree species are known to vary with rising temperatures (Berry and Bjorkman 1980), consequently influencing the potential carbon uptake of trees under future climates. Previous studies have indicated that the impacts of temperature variability on photosynthesis contribute considerably to the uncertainty in the terrestrial carbon cycle (Booth et al. 2013, Mercado et al. 2018) and climate change prediction (Friedlingstein et al. 2006). Therefore, advancing the understanding of the temperature dependence of photosynthesis is critical to constrain the uncertainty in vegetation carbon uptake, and thus critical to evaluate the effectiveness of reforestation and afforestation for mitigating climate change.

Previous studies have explored the temperature dependence of key photosynthetic traits (i.e., the maximum carboxylation rate, V_cmax_, and the maximum electron transport rate, J_max_) for species across the globe (Ainsworth and Long 2005, Slot and Winter 2017). While some were able to develop a unified function that describes the temperature dependence across species (Medlyn, Dreyer, et al. 2002, Medlyn, Loustau, et al. 2002), other studies found significant inter-species variation in the magnitude and shape of the temperature dependence (Way and Oren 2010). This temperature dependence of V_cmax_ and J_max_ based on limited and divergent observations is then implemented in Dynamic Global Vegetation Model (DGVMs) or Earth System Models (ESMs) for large-scale simulation of the carbon cycle (Mercado et al. 2018). However, these previous studies greatly lack the observations in tropical Asia (Crous et al. 2022). Tropical Asia hosts some of the most carbon-dense (Saatchi et al. 2011) and diverse ecosystems (Slik et al. 2015) in the world. Currently, the temperature dependence of V_cmax_ and J_max_ for species in tropical Asia is extrapolated from either other parts of the tropics (e.g., French Guiana, Coste et al. (2005)) or the high-temperature scenarios for extra-tropics species (Slot and Winter 2017). A recent study demonstrated that Southeast Asia has the most considerable uncertainty in GPP across models (Bastos et al. 2020), further emphasizing the lack of observational constraints for temperature dependence of photosynthesis in tropical Asia.

Therefore, in this study, we aimed to study the temperature dependence of photosynthesis for the common tree species in Singapore, a city-state in tropical Southeast Asia. We selected the nine most common species planted in Singapore’s past and present extensive reforestation efforts, namely the Garden City Initiative and the One Million Tree movement. These species are also widely distributed across Southeast Asia. Our research objectives are to: 1) quantify the V_cmax_ and J_max_ of across tree species in tropical Asia; 2) determine how V_cmax_ and J_max_ respond to temperature for the tree species and 3) explore the physiological mechanism for the inter-species variation in the temperature dependence of photosynthesis for these species.

## Materials and methods

### Study site and tree species

We conducted our fieldwork from July 2023 to January 2024 across Singapore, which is located near the equator (1.283°N, 103.833°E) and is within the intertropical convergence zone. Singapore’s annual temperatures average 27–30ºC, with a daily mean maximum of 30–31ºC. These temperatures have increased by 0.25ºC per decade from 1948 to 2023 (Marzin et al. 2015). The annual total precipitation hovers around 2800 mm year^−1^ with an average relative humidity of *c*.*a*. 80% (Acero et al. 2024).

We sampled the nine most common tree species planted in Singapore (Fig. 1; Table S1). We selected these tree species based on the records from Singapore’s reforestation initiatives, such as the One Million Tree Movement and the Garden City Campaign. These tree species are native to either Singapore or Southeast Asia or have been introduced to Singapore for more than a century. In particular, we identified the representative individuals of these tree species in Bishan-Ang Mo Kio Park (1.3634°N, 103.8436°E), Singapore Botanic Gardens (1.3138°N, 103.8159°E), Jurong Lake Gardens (1.3359°N, 103.7262°E), East Coast Park (1.3008°N, 103.9122°E) and Pasir Ris Mangrove Boardwalk (1.3715°N, 103.9524°E).

**Figure 1.**
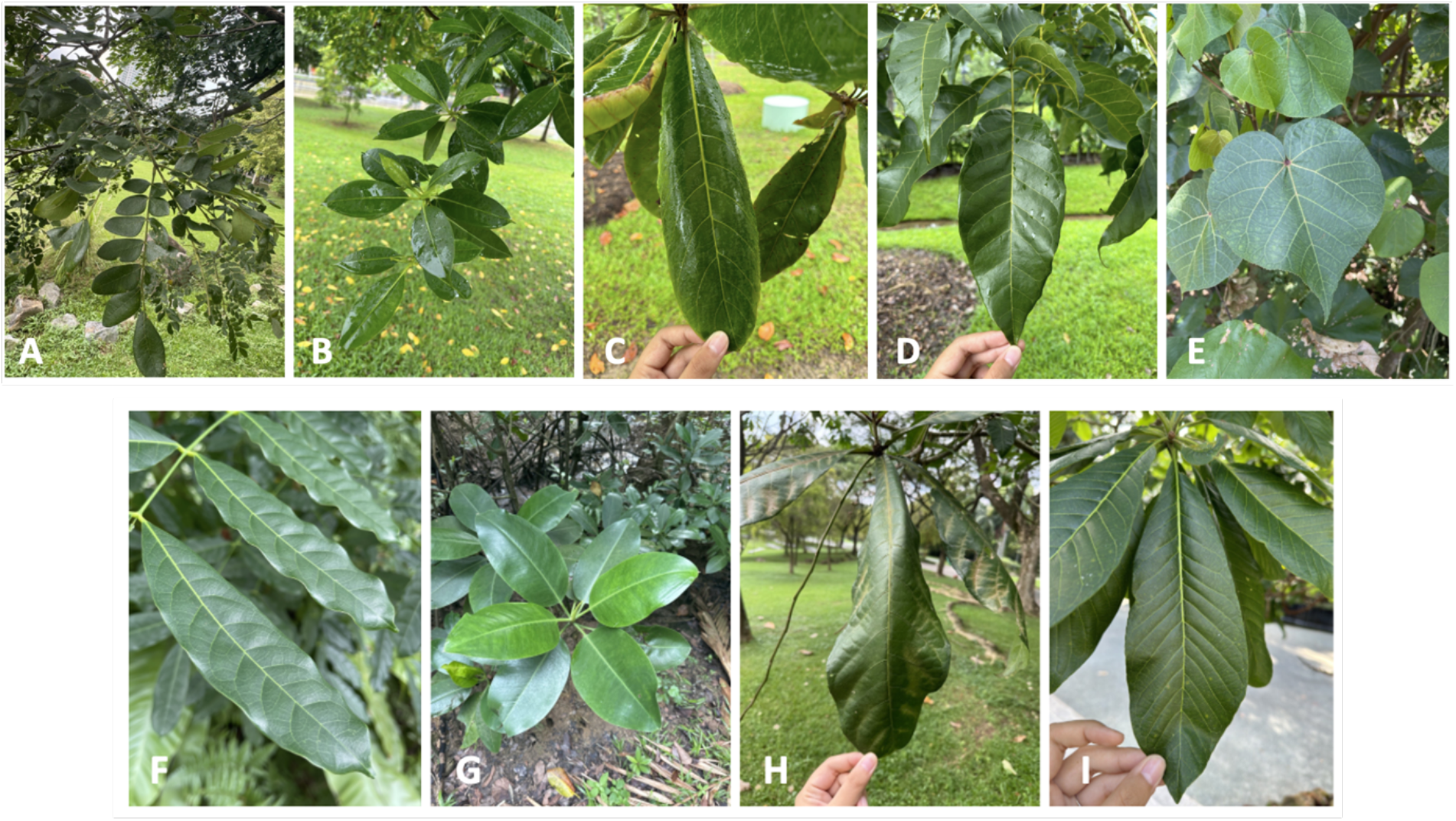
Images of the sampled trees species: (A) Rain Tree (*Samanea saman*), (B) Tembusu (*Cyrtophyllum fragrans*), (C) Sea Almond (*Terminalia catappa*), (D) Pink Poui (*Tabebuia rosea*), (E) Sea Hibiscus (*Hibiscus tiliaceus*), (F) African Mahogany (*Khaya senegalensis*), (G) Tall-stilt Mangrove (*Rhizophora apiculata*), (H) Fish Poison (*Barringtonia asiatica*), (I) Cannonball Tree (*Couroupita guianensis*).

### Gas exchange measurements and curve fitting

In this study, we used the LICOR 6800 Portable Gas Exchange System (Li-Cor Inc., 2019) to measure the CO_2_ dependence of leaf photosynthesis (i.e., A-C_i_ curve) under a temperature gradient from 30 to 34ºC. Three healthy and mature sunlit leaves per species were selected. Then five A-C_i_ curves of each leaf were recorded at every 1ºC interval from 30ºC to 34ºC. The CO_2_ concentrations in the reference chamber for the A-C_i_ curves were set in a sequence of 400, 300, 200, 100, 50, 0, 400, 400, 600, 800, 1000, 1200, and 1600 μmol mol^−1^. We set a saturating light intensity at 1600 μmol m^−2^ s^−1^ and kept the relative humidity in the leaf chamber greater than 30% and less than 80%. We then fitted the A-C_i_ curves using the R package “*plantecophys*” (Duursma 2015) and extracted V_cmax_ and J_max_. This package employs the Farquhar, von Caemmerer, and Berry (FvCB) photosynthesis model (Farquhar et al. 1980).

### Estimation of leaf photosynthesis along a warming gradient

To compare the temperature dependence of photosynthesis of these common species against the temperature-dependence prescribed in Earth System Models, we estimated the temperature dependence of photosynthesis using the model from Medlyn, Loustau, et al (2002) and Kattge and Knorr (2007), which has been adopted in Earth System Models (e.g., the CABLE model, (Haverd et al. 2018) and the CLM model, Smith et al (2017)). We first employed the global map of V_cmax_ at 25°C (Luo et al. 2021, spatial resolution: 0.5°×0.5°). We then extracted Singapore’s V_cmax_ at 25°C using its longitude and latitude. After that we estimated the V_cmax_ from 30 to 34ºC using a peaked equation as follows:

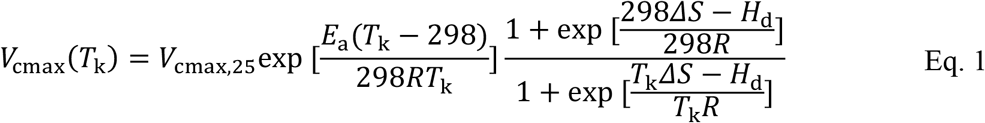

in which *V*_cmax_(*T*_k_) denotes V_cmax_ at a certain temperature (*T*_k_, in K), *V*_cmax, 25_ is the value of V_cmax_ or J_max_ at 25°C, *E*_a_ and *H*_d_ represent the exponential rate of the increase and decrease of the function (71500 and 20000 J mol^−1^ respectively), *R* is the universal gas constant (8.314 J mol^−1^ K^−1^), and *Δ*S is an entropy factor (650 J mol^−1^ K^−1^). We then estimated J_max_ using its empirical relationship to V_cmax_ under varying temperatures (*T*, in °C) according to Leuning (2002):

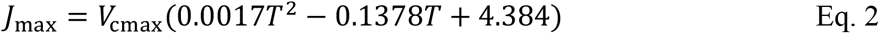

We then input the values of V_cmax_ and J_max_ into the FvCB model and estimated A_n_ using the “*plantecophys*” package. Here we assumed that atmospheric CO_2_ concentration is 400 μmol mol^−1^, vapor pressure deficit is 0.75 kPa, and light intensity is 1200 μmol m^−2^ s^−1^.

## Results

### Species-specific temperature dependence of V_cmax_ and J_max_

We found that V_cmax_ and J_max_ across species vary substantially in magnitude as well as in their temperature dependence (Fig. 2). Among the 9 species, Sea Hibiscus had the largest V_cmax_ (56 μmol m^−2^ s^−1^), followed by Rain Tree, Sea Almond, and African Mahogany (from 40 to 50 μmol m^−2^ s^−1^) while the remaining species (Cannonball, Fish Poison, Pink Poui, Tall-stilt Mangrove, and Tembusu) had lower V_cmax_ *c*.*a*. 20–30 μmol m^−2^ s^−1^. Sea Almond also had the largest J_max_ of 110 μmol m^−2^ s^−1^, followed by Rain Tree and Sea Hibiscus (100–120 μmol m^−2^ s^−1^), while J_max_ of the other species varied from 40 to 80 μmol m^−2^ s^−1^.

**Figure 2.**
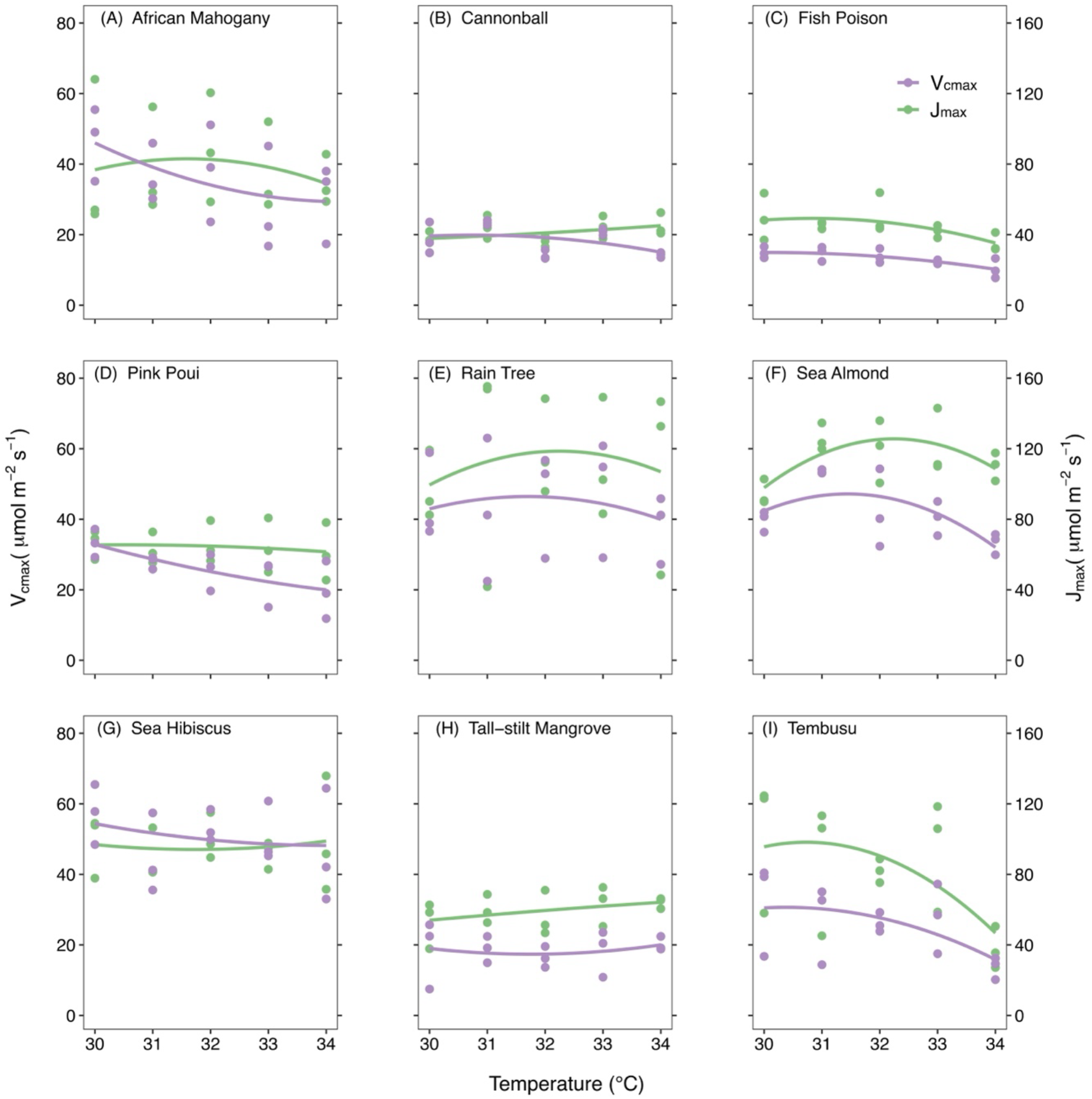
The temperature dependence of maximum carboxylation rate (V_cmax_) and maximum electron transport rate (J_max_) of the ten common tree species: (A) Rain Tree, (B) Tembusu, (C) Sea Almond, (D) Pink Poui, (E) Sea Hibiscus, (F) African Mahogany, (G) Tall-stilt Mangrove, (H) Fish Poison, (I) Cannonball Tree. We show the fitted quadratic temperature responses of V_cmax_ (purple) and J_max_ (green).

Furthermore, we generally observed a decline of V_cmax_ with rising temperature (six out of the nine species), with the most significant declines observed for African Mahogany and Tembusu (Fig. 2a, i). Over the 5ºC gradient from 30 to 34 ºC, V_cmax_ declined by nearly 50%. Rain Tree and Sea Almond demonstrate an optimal temperature around 31–32ºC. In contrast, we found that the V_cmax_ of Cannonball, Fish Poison, Sea Hibiscus, and Tall-stilt Mangroves is not sensitive to temperature.

We observed slightly different patterns of J_max_ change with rising temperature. Five out of the nine species demonstrated increase of J_max_ with temperature, with the most significant increase observed for Tall-stilt Mangroves and Sea Almond (20–30%). Tembusu is the only species showing a decrease of J_max_ with rising temperature. African Mahogany, Rain Tree, and Sea Almond demonstrated an optimal temperature of around 31–32ºC. The J_max_ of Cannonball, Pink Poui, and Sea Hibiscus was not sensitive to temperature.

### Differing temperature responses of net photosynthetic rate

We then estimated the net photosynthetic rate (A_n_) of nine species over a 5°C temperature gradient using the observed V_cmax_ and J_max_ in the FvCB model (see methods). Rain Tree and Sea Hibiscus showed generally higher A_n_ than that of Sea Almond, Tembusu, Pink Poui, and African Mahogany, while Cannonball, Tall-stilt Mangrove, and Fish Poison showed the lowest A_n_ (Fig. 3). Despite the differing levels of A_n_ across species, we also found that the patterns of temperature responses of A_n_ considerably varied across species. Rain Tree, Sea Almond, and Tembusu exhibited a maximum A_n_ under 31–32°C (Fig. 3). A_n_ of Pink Poui and African Mahogany declined with temperature, while A_n_ of Sea Hibiscus, Cannonball, Tall-stilt Mangrove, and Fish Poison did not show clear changes to temperature. Furthermore, we estimated the theoretical A_n_ under Singapore’s climate using the local V_cmax_ at 25°C provided in (Luo et al. 2021) and the models provided in Medlyn, Loustau, et al. (2002) and Leuning (2002) (see Methods). We found that the A_n_ estimated at the local species level were generally lower than the estimated A_n_ (Fig. 3, black triangles) from the global datasets and models.

**Figure 3.**
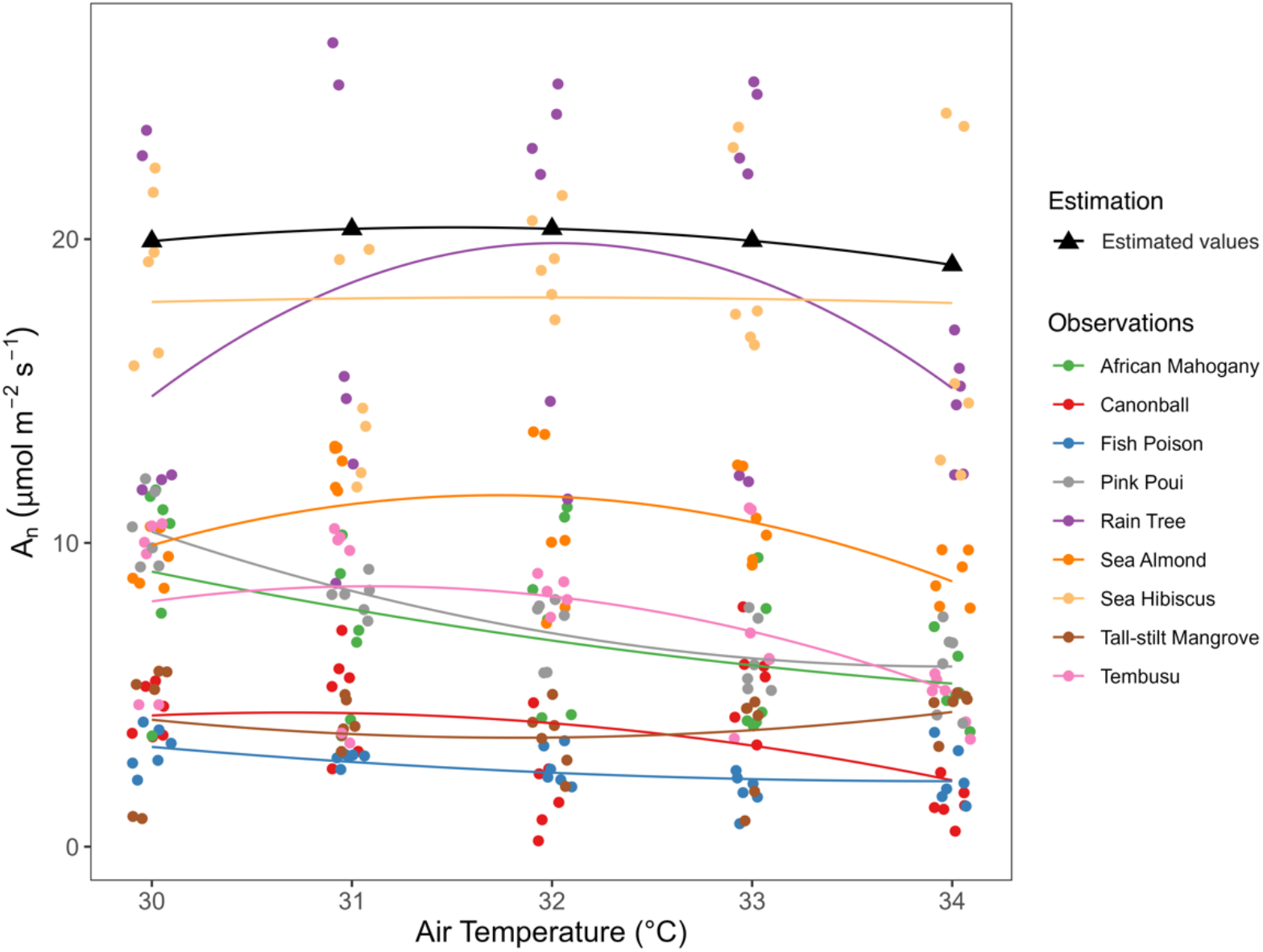
The responses in net photosynthetic rate (A_n_) to a range of air temperatures (30– 34°C). Each dot indicates one observation of A_n_ and the palette shows observations from different species. The black dots denote the estimated theoretical A_n_ under the climate of Singapore (see Methods). Solid lines were fitted using quadratic equations.

### Leaf thermal regulation coordinates with temperature sensitivity of V_cmax_ and J_max_

We further explored the inter-species variations in the temperature dependence of V_cmax_ and J_max_ and examined their relationship with leaf thermal regulation. The temperature dependence of V_cmax_ and J_max_ were indicated by the coefficient of variation (CV) for V_cmax_ and J_max_ under different temperatures – if the CV was greater, it indicates a greater temperature sensitivity. We found the CV of V_cmax_ varies substantially across species, from less than10% (e.g., Sea Hibiscus, Rain Tree, and Tall-stilt Mangrove) to above 30% (e.g., Tembusu) (Fig. 4a). The CV of V_cmax_ was generally greater than that of J_max_ except for Tembusu (Fig. 4b). We then calculated the difference of leaf temperature to air temperature (T_diff_), as an indicator of leaf thermal regulation. Since in the tropical daytime, leaf temperature is often higher than air temperature, therefore a smaller T_diff_ indicate a greater thermal regulation. We found that the species with a small T_diff_ tend to exhibit a low CV of V_cmax_ and J_max_ (e.g., Sea Hibiscus) and *vice versa* (e.g., Tembusu), although the correlations between CV and T_diff_ were only close to be significant (V_cmax_: r = 0.48, *p* = 0.20; J_max_: r = 0.58, *p* = 0.10).

**Figure 4.**
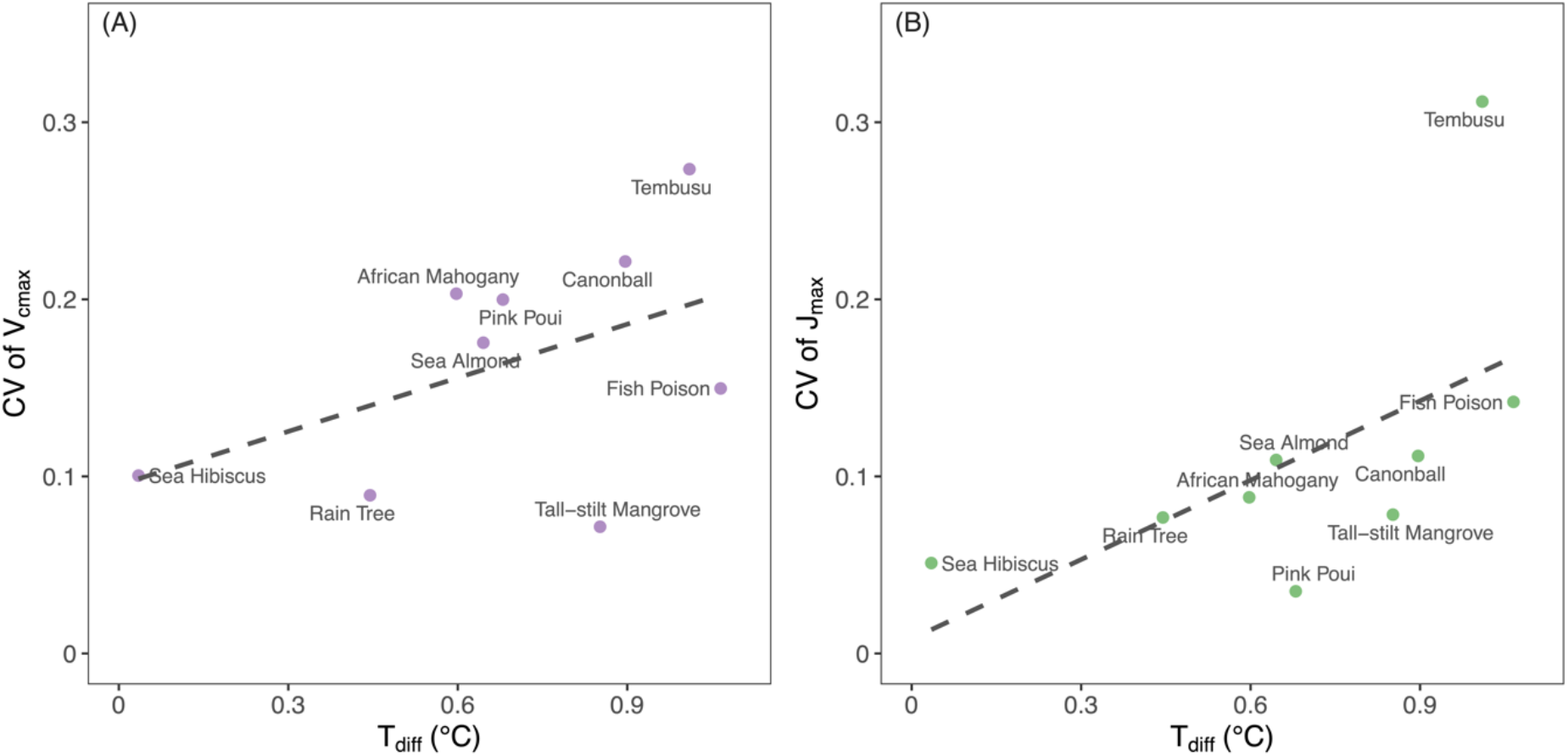
The coordination between the thermal variability of photosynthetic capacity (coefficient of variations, or CV of V_cmax_ (A) and J_max_ (B)) and the difference between leaf temperature and air temperature (T_diff_). CV was calculated using values of V_cmax_ and J_max_ under five air temperatures (30–34°C). Tdiff was calculated as T_leaf_−T_air_. The dashed line indicates the correlation between CV and T_diff_.

## Discussion

In this study, we found substantial variations (i.e., greater than three-fold) in photosynthetic capacity (V_cmax_ and J_max_) among nine common tree species in the tropical state of Singapore. There was a large discrepancy in their temperature dependence, where we found the photosynthesis of Cannonball, Fish Poison, Sea Hibiscus, and Tall-stilt Mangroves are not very sensitive to rising temperature. Species that better adjust leaf temperature according to air temperature (with a small T_diff_) tend to exhibit low temperature dependence of their photosynthetic capacity, suggesting that the capacity of leaf thermal regulation contributes to photosynthetic thermal variability. These results provide valuable information on the photosynthesis dynamics of understudied tree species in tropical Southeast Asia.

### Inter-species variation in photosynthetic capacity

In this study, we found considerable variations in the temperature response of photosynthesis among the common tree species. We could not explain these variations by the origin of the species, leaf lifespan, or tree height (Table S1). For instance, Rain Tree and Cannonball share similar leaf lifespan and tree height, however, they differed considerably in their photosynthetic rates and CV of V_cmax_. Moreover, studies have found that coexisting species under similar climate exhibit significant variations in photosynthetic traits (e.g., A_max_ and leaf mass per area), suggesting distinctly different functional modulations among species irrespective of biomes or climate (Wright et al. 2004, Wang et al. 2022). Therefore, our study emphasizes that enhanced observations of photosynthetic traits of more representative species are necessary for constraining the uncertainty in carbon modeling of tropical Asia.

Nevertheless, we noticed that tree species mostly planted in Garden City Initiative (launched in 1967), such as Rain Tree, Sea Hibiscus, Sea Almond, and Tembusu that we observed, had higher A_n_ than those planted in the more recent One Million Tree Movement (launched in 2020), such as Fish Poison and Tall-stilt Mangrove. This suggests that leaves from mature trees likely photosynthesized more than those from younger trees. Although we were not able to obtain the extract age of the individual trees, the trees planted in the One Million Tree Movement were likely less than 3 years old. Currently the impacts of stand demography on leaf photosynthesis remain unclear (Bond 2000), as some suggested that the higher photosynthesis of older trees might benefit from deeper roots, thus facilitating the absorption of water and nutrition in dry seasons. There are some evidence indeed suggested that mature oil palm trees tend to have greater photosynthesis capacity than younger trees in Southeast Asia (Apichatmeta et al. 2017) Meanwhile, the impact of leaf demography (i.e., young and mature leaves) on photosynthetic capacity has been widely recognized in the tropics (Wu et al. 2016, Albert et al. 2018) and adopted in recent modelling efforts (Manoli et al. 2018). However, since we collected most of our data in the late dry season, we assumed limited impacts from leaf demography in our analysis.

In addition, our result suggests that empirical temperature response model overestimated A_n_ compared to observed A_n_ of all species in our sties and underestimated the temperature variability of A_n_ (Fig. 3). This could be attributed to the fact that values of the parameters for temperature dependence were mostly extrapolated from temperature or boreal plant species (e.g., *Nicotiana tabacum* and *Abies alba*), thus not reflecting the temperature sensitivity of photosynthesis for tropical tree species. Moreover, temperature regulates stomatal conductance, directly limiting photosynthesis more under increasing temperature. However, the altered stomatal conductance influence leaf thermal properties, thus exerting biochemical limitation to photosynthetic apparatus (Slot and Winter 2017). This indirect effect is not reflected by the biochemical model alone (Bambach et al. 2020). Therefore, a better understanding of linking the role of physiological acclimation to stomatal dynamics is needed for improved predictions of carbon assimilation under warming climate.

### Inter-species variation in temperature dependence of photosynthesis and its coordination with leaf thermal adjustment

In this study, we found that the CV of V_cmax_ and J_max_ was positively correlated to the T_diff_, indicating the potential role leaf thermal regulation capacity on the inter-species variation of leaf photosynthetic temperature dependence. The large spread in CV and T_diff_ suggest divergent strategies of plants to coordinate thermal regulation and photosynthetic variability. In particular, species with a smaller T_diff_ (i.e., stronger capacity to transpire water and reduce leaf temperature) tends to have low CV of V_cmax_ and J_max_, thus leading to a more stable photosynthetic performance. The capacity to maintain a small CV is related to the plasticity of stomatal conductance (G_s_) under different temperatures, as the Sea Hibiscus had a much larger plasticity of G_s_ than Tembusu and Fish Poison (Fig. 5). Although the inter- or intra-species variations of photosynthetic thermal dependence have been explored in previous studies (Zaka et al. 2016, Vico et al. 2019), our work clearly related these variations to the stomatal behaviour and the capacity of leaf to regulate its temperature.

**Figure 5.**
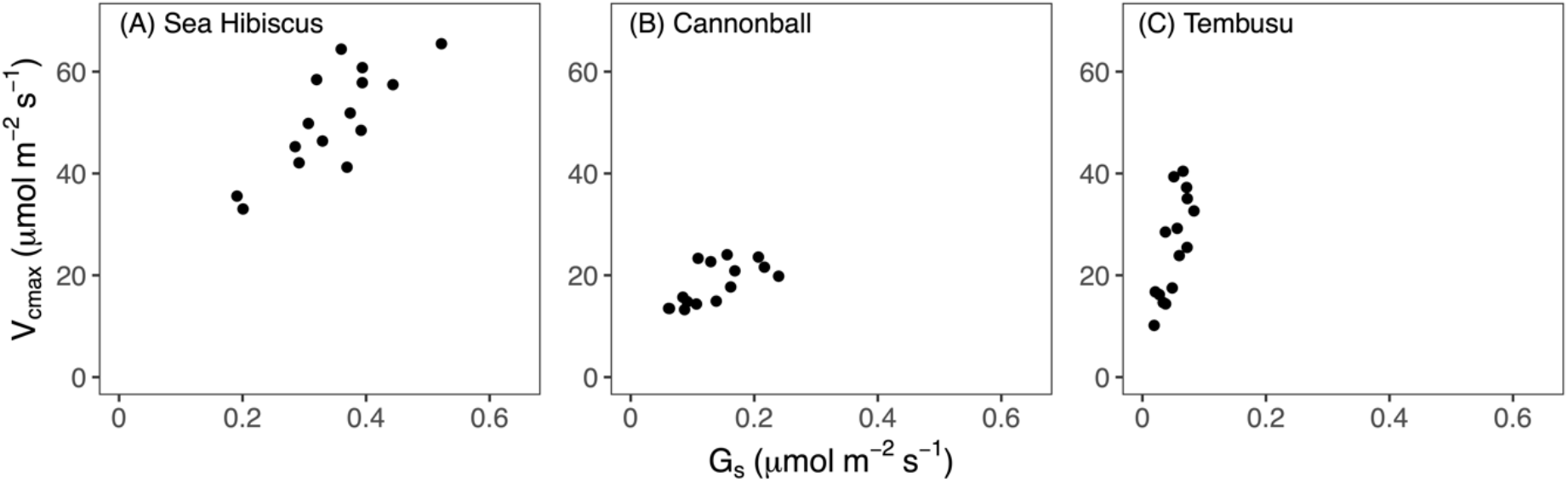
The correlation between V_cmax_ and stomatal conductance (G_s_) of species differing in the plasticity of G_s_: (A) Sea Hibiscus, (B) Cannonball, and (C) Tembusu.

### Implication for guiding tree planting initiatives

Our study provides the critical information to select the suitable tree species for tree-planting initiatives aimed at climate change mitigation. This is especially relevant to Singapore due to its land scarcity and profound urban heat island effect (Grace Wong 2004, Chow and Roth 2006). Therefore, selecting tree species that have maximized and sustained photosynthesis under higher temperatures will help ensure the effectiveness of reforestation initiatives in the future. We found that Sea Hibiscus and Rain Tree tend to have high A_n_, but only Sea Hibiscus has low thermal variability under future rising temperatures. Pink Poui and African Mahogany show clear declines in A_n_ when temperature rises. Sea Almond and Tembusu, along with Rain Tree have an optimal temperature of photosynthesis around 32°C, beyond which their photosynthesis would decline. Cannonball, Fish Poison, and Tall-stilt Mangroves show high degree of temperature resistance in their photosynthesis, though their photosynthesis rate is lower than other major tree species. Overall, Sea Hibiscus seems to be the only species that has great and sustained carbon sequestration potential under future climate than other species. However, we would also like to acknowledge that our reported dependence here are based on short-term temperature responses, and over long-term the acclimation of photosynthesis to temperature (Medlyn, Dreyer, et al. 2002, Kattge and Knorr 2007) may impact the inter-species variation we observed. Nevertheless, our result provides a much needed set of observations on the photosynthesis of common species in tropical Asia, and will guide the reforestation efforts in Singapore and Southeast Asia.

## Conclusion

In summary, our study revealed substantial variations in the temperature responses of photosynthesis among nine tree species commonly planted in Singapore’s tree planting initiatives. Our study underscores the need for a strategic approach to tree selection in these initiatives, considering temperature dependence of photosynthesis, to optimise the carbon sequestration potential of urban forests and mitigate climate change. Our study also demonstrates that inter-species difference in the temperature dependence of photosynthesis is likely linked to the capacity of leaf thermal regulation of each species. This finding deepens our understanding of the physiological mechanisms underlying photosynthetic temperature dependence and facilitates the identification of tree species that are resilient to future climate.

## Acknowledgments

The authors would like to thank National Biodiversity Centre, National Parks Board of Singapore for providing the license for fieldwork in Singapore Botanical Gardens.

## Authors’ contributions

L.M.R.T. and X.L. conceived and designed the study; X.L. secured the resources and fund for the study; L.M.R.T. performed data collection and analysed the data; L.M.R.T. and L.Y. led the writing of the manuscript; all authors contributed critically to the writing and revision of the manuscript and gave final approval for the submission.

## Supplementary data

Supplementary data are available in a separate document.

## Data availability

The data collected and presented in the study will be uploaded to the TRY Plant Trait Database (https://www.try-db.org/TryWeb/Home.php) upon the acceptance of the manuscript.

