## Supplemental Table 1 for "Variations in the temperature dependence of photosynthesis among nine common tree species planted in Singapore"

**Supplementary materials**

| **Table S1**. Summary of the tree species investigated in this study. | | | | | | | |
| --- | --- | --- | --- | --- | --- | --- | --- |
| Species | Family | Origin | Height | Leaf lifespan | Preferred climatic zone | Native habitat | Singapore restoration initiative |
| *Rain Tree* | Fabaceae | Tropical America | Up to 30 m | 0.78–1 year | Tropical | Terrestrial (Primary Rainforest, Grassland / Savannah/ Scrubland) | Garden City initiative |
| *Tembusu* | Verbenaceae | Southeast Asia | Up to 30 m | ~0.78 year | Tropical, Sub-Tropical / Monsoonal | Terrestrial (Primary Rainforest, Secondary Rainforest, Monsoon Forest, Coastal Forest, Freshwater Swamp Forest, Disturbed Area / Open Ground) | Garden City initiative, Heritage Tree |
| *Sea Almond* | Combretaceae | Tropical Asia | Up to 35 m | >1 year | Tropical, Sub-Tropical / Monsoonal | Terrestrial (Coastal Forest), Shoreline (Mangrove Forest, Sandy Beach, Rocky Beach) | Garden City initiative |
| *Pink Poui* | Bignoniaceae | Mexico | Up to 35 m | 0.78–1 year | Tropical | Terrestrial (Primary Rainforest, Secondary Rainforest) | Garden City Initiative |
| *Sea Hibiscus* | Meliaceae | Tropical Asia, Americas | Up to 30 m | >1 year | Tropical | Terrestrial (Coastal Forest), Shoreline (Mangrove Forest, Sandy Beach) |  |
| *African Mahogany* | Rhizophoraceae | Tropical West Africa | Up to 20 m | 0.78–1 year | Tropical Savanna | Riverine forests | Garden City Initiative, One Million Tree Movement |
| *Tall-stilt Mangrove* | Lecythidaceae | Singapore | Up to 30 m | >1 year | Tropical, Sub-Tropical / Monsoonal | Shoreline (Mangrove Forest) | One Million Tree Movement, Mangrove Restoration |
| *Fish Poison* | Malvaceae | Singapore | Up to 15 m | >1 year | Tropical | Terrestrial (Coastal Forest) | One Million Tree Movement |
| *Cannonball* | Lecythidaceae | Tropical South America | Up to 35 m | 0.78–1 year | Tropical | Terrestrial (Primary Rainforest) | Garden City Initiative, One Million Tree Movement, Heritage Trees |
